# CryoEM structure of the EBV ribonucleotide reductase BORF2 and mechanism of APOBEC3B inhibition

**DOI:** 10.1101/2021.08.30.458246

**Authors:** Nadine M. Shaban, Rui Yan, Ke Shi, Sofia N. Moraes, Adam Z. Cheng, Michael A. Carpenter, Jason S. McLellan, Zhiheng Yu, Reuben S. Harris

## Abstract

Viruses use a plethora of mechanisms to evade immune responses. A new example is neutralization of the nuclear DNA cytosine deaminase APOBEC3B by the Epstein-Barr virus (EBV) ribonucleotide reductase subunit BORF2. Cryo-EM studies of APOBEC3B-BORF2 complexes reveal a large >1000 Å^2^ binding surface comprised of multiple structural elements from each protein, which effectively blocks the APOBEC3B active site from accessing single-stranded DNA substrates. Evolutionary optimization is suggested by unique insertions in BORF2 absent from other ribonucleotide reductases and preferential binding to APOBEC3B relative to the highly related APOBEC3A and APOBEC3G enzymes. An atomic understanding of this novel pathogen-host interaction may contribute to the development of drugs that block the interaction and liberate the natural antiviral activity of APOBEC3B.

**One-Sentence Summary:** These studies show how a conserved viral nucleotide metabolism protein is repurposed to inhibit a potent antiviral factor.

---

Ribonucleotide reductases (RNRs) are indispensable for DNA-based organisms by catalyzing the conversion of ribonucleotides to deoxyribonucleotides [reviewed by (*1*)]. This fundamental activity makes RNRs targets for anticancer, antibacterial, and antiviral therapies [reviewed by (*1*)]. Viruses augment deoxyribonucleotide supplies by altering pathways that regulate RNR levels in the host, infecting only replicating cells that would have high levels of RNR activity or, in the case of many large DNA viruses and bacteriophages, encoding their own RNRs [reviewed by (*2, 3*)]. Herpesvirus RNRs are similar to those in humans, yeast, and some bacteria, and belong to the class Ia subgroup consisting of a large and small subunit. These RNRs are regulated allosterically by ATP/dATP, and by intricate oligomerization mechanisms that ensure a balanced supply of each nucleotide precursor (*1, 4, 5*). However, it has been unclear if viral RNRs utilize the same regulatory and catalytic mechanisms with some evidence suggesting differences. For example, viral RNRs are unresponsive to dATP and other effectors that regulate activity (*6, 7*), β-herpesvirus RNRs lack a gene that encodes a small subunit and thus are presumed inactive (*2, 3*), and HSV-1 RNRs have n-terminal extensions that interact with cellular factors unrelated to RNR catalytic activity (*2, 3*). More recently, we discovered that the large subunit of the Epstein-Barr virus (EBV) RNR, BORF2, interacts with at least two cellular APOBEC3 enzymes revealing an unexpected role for herpesvirus RNRs in blocking antiviral innate immunity (*8, 9*).

The seven-membered APOBEC3 family (A3A-D, F-H) of single-stranded DNA cytosine deaminases elicits broad antiviral activity, often resulting in lethal mutagenesis, with the archetypical example being HIV-1 hypermutation (counteracted by the HIV-1 Vif protein that targets restrictive A3s for polyubiquitination and degradation) [reviewed by (*10, 11*)]. In recent years, A3 mutagenesis has also become synonymous with cancer as dysregulation of A3A/A3B is linked to heavy mutational burdens found in bladder, breast, cervical, head/neck, lung, and other tumor types [reviewed by (*12, 13*)]. Interestingly, EBV BORF2, and related RNRs from KSHV1 and HSV-1 are capable of relocalizing these two deaminases from the nuclear compartment into cytoplasmic aggregates, most likely to protect viral genomes from deamination during the lytic phase of DNA replication (*8, 9, 14*). In line with this, BORF2 potently inhibits the DNA deaminase activity of A3B (*8*). To date, no eukaryotic viral RNR structures have been determined and the structural basis for the RNR interaction with A3 enzymes is unknown. The primary aim of the present studies is to determine the structural basis and mechanism for this unanticipated host-pathogen interaction.

Prior studies indicated that the BORF2-A3B interaction minimally requires the BORF2 RNR core domain and the A3B c-terminal catalytic domain (ctd) (*8*). We therefore focused on forming a complex between BORF2 and A3Bctd. Initial attempts to purify BORF2 from *E. coli* were unsuccessful. However, expression of BORF2 in 293T cells with an n-terminal maltose binding protein (MBP) tag significantly improved solubility, purity, and yield (**Fig. 1A**). Catalytically active A3Bctd with a c-terminal mychis tag was also expressed in 293T (**Fig. 1A**). The purified BORF2 protein efficiently inhibits A3B catalytic activity *in vitro* (**Fig. 1B**). To form the BORF2-A3B complex, BORF2 and A3Bctd were purified individually with affinity resins then mixed together with excess A3Bctd and the complex was isolated by size-exclusion chromatography (**Fig. 1A**). This complex migrated at a size consistent with a higher-order assembly and a 1:1 ratio of BORF2 to A3B (**Fig. 1A**).

**Fig. 1.**
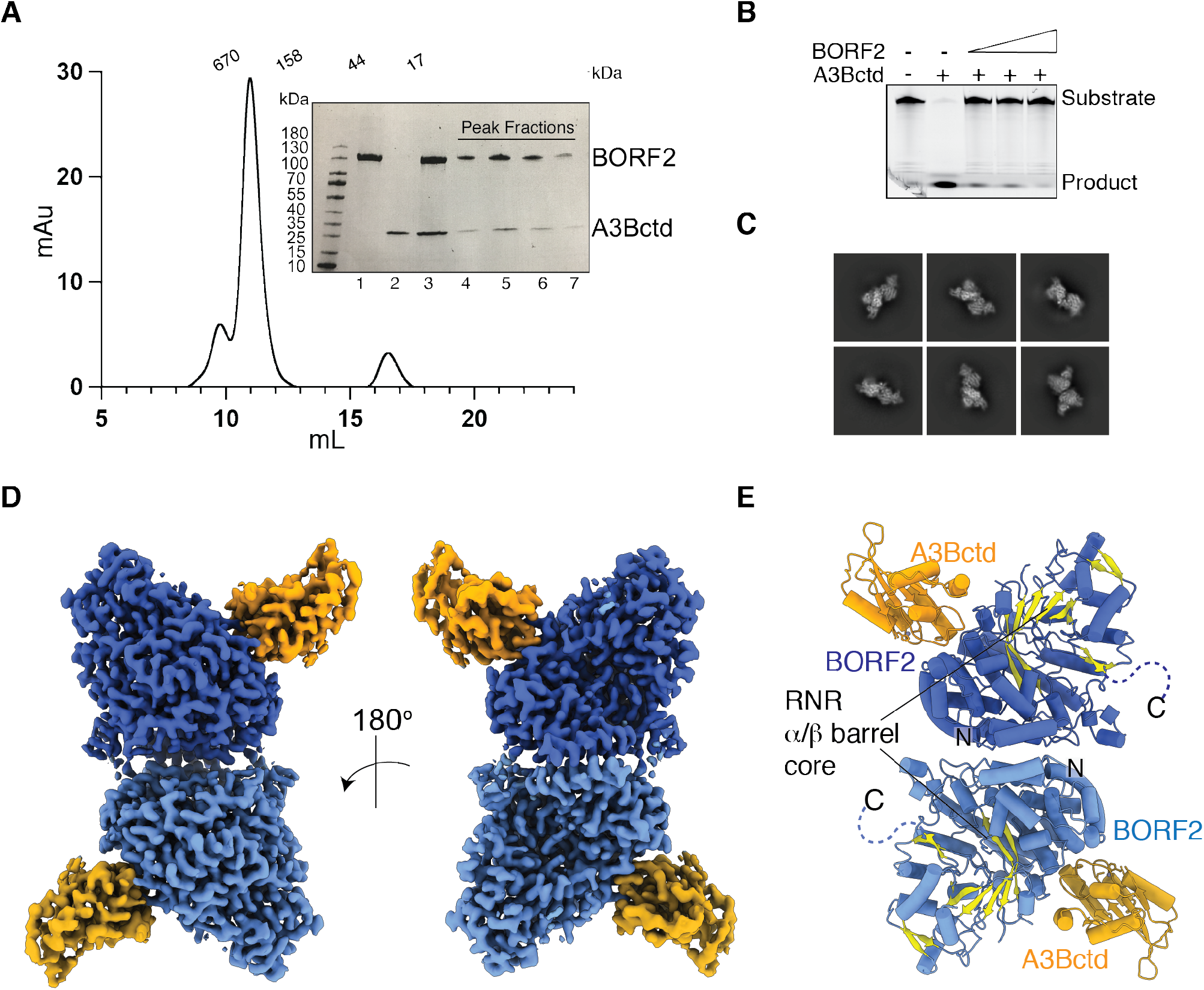
Structure of EBV BORF2-A3Bctd complex. **(A)** Size exclusion chromatogram for BORF2-A3Bctd complex. Inset: SDS page gel of MBP-BORF2, A3Bctd-mychis, the complex prior to SEC (lanes 1-3), and fractions of the major peak (lanes 4-8). **(B)** BORF2 inhibits A3Bctd ssDNA deaminase activity. Negative control is substrate alone, positive control is A3Bctd alone, and dose responsive inhibition occurs with increasing BORF2 concentrations. **(C)** Representative 2D classes of the BORF2-A3Bctd complex. **(D)** Cryo-EM composite map of BORF2-A3Bctd complex with BORF2 in blue and A3Bctd in orange. **(E)** Ribbon schematic of the BORF2-A3Bctd complex with the RNR core depicted in yellow.

Cryoelectron microscopy (cryo-EM) was used to determine the structure of the BORF2-A3Bctd complex. A total of 7,488 multi-frame movies were collected using a Titan Krios with a post energy filter K3 detector and processed in cryoSPARC (*15*). 2D classes corresponding to what appeared to be dimeric forms of the complex were selected for 3D reconstruction and heterogeneous refinement yielding a 2.8 Å map. 3D variability analysis was performed to assess the conformational flexibility of the complex (*16*). Movement was particularly detected at the BORF2 dimer interface (movies S1 and S2). Therefore, a focused refinement around the BORF2-A3B monomer was performed after symmetry expansion that resulted in an improved reconstruction to 2.55 Å. A composite map to represent the dimeric form of the complex was generated from the focused refined map (fig. S1-2, table S1). The cryo-EM maps allowed the modelling of nearly all components of the BORF2-A3Bctd complex with higher resolution features at the BORF2 catalytic core and between the BORF2-A3B interface (fig. S2). The structure is composed of a dimer of BORF2 monomers, which are each bound to one A3Bctd protomer (**Fig. 1C-E**).

BORF2 adopts an α/β barrel RNR architecture resembling the large subunit of all classes of RNRs (*17*) (**Fig. 1E**, fig. S3). Although sharing only ∼25% primary amino acid identity, the structure of BORF2 overlays well with the RNRs of *E. coli* and humans with r.m.s.d. of 1.1, and 1.2 Å, respectively (*e*.*g*., **Fig. 2A**, fig. S3A). Particularly high conservation is observed for residues involved in ribonucleotide reduction (fig. S3B-C). For class I RNRs, ribonucleotide reduction involves the transfer of a radical generated in the iron coordinating small subunit of the RNR to the large subunit via tyrosines and cysteine amino-acid radical intermediates (fig. S3B) (*1*). The resulting thiyl radical formed in the large subunit initiates the reduction of the 2-OH of the ribonucleotide and subsequent oxidation of a pair of redox active cysteines positioned on adjacent strands. BORF2 residues Y725, Y726, C389, C187, and C403 overlay with the established catalytic residues of other class Ia RNRs involved in this process (fig. S3C). The predicted redox active cysteines (C187, C483) are in a reduced state in the BORF2 structure. The c-terminal end of BORF2 that is predicted to contain additional cysteines required for post-catalytic re-reduction of the active site cysteines is disordered similar to other class Ia RNRs [*e*.*g*., (*18-20*); dashed line to c-terminus in **Fig. 1E**].

**Fig. 2.**
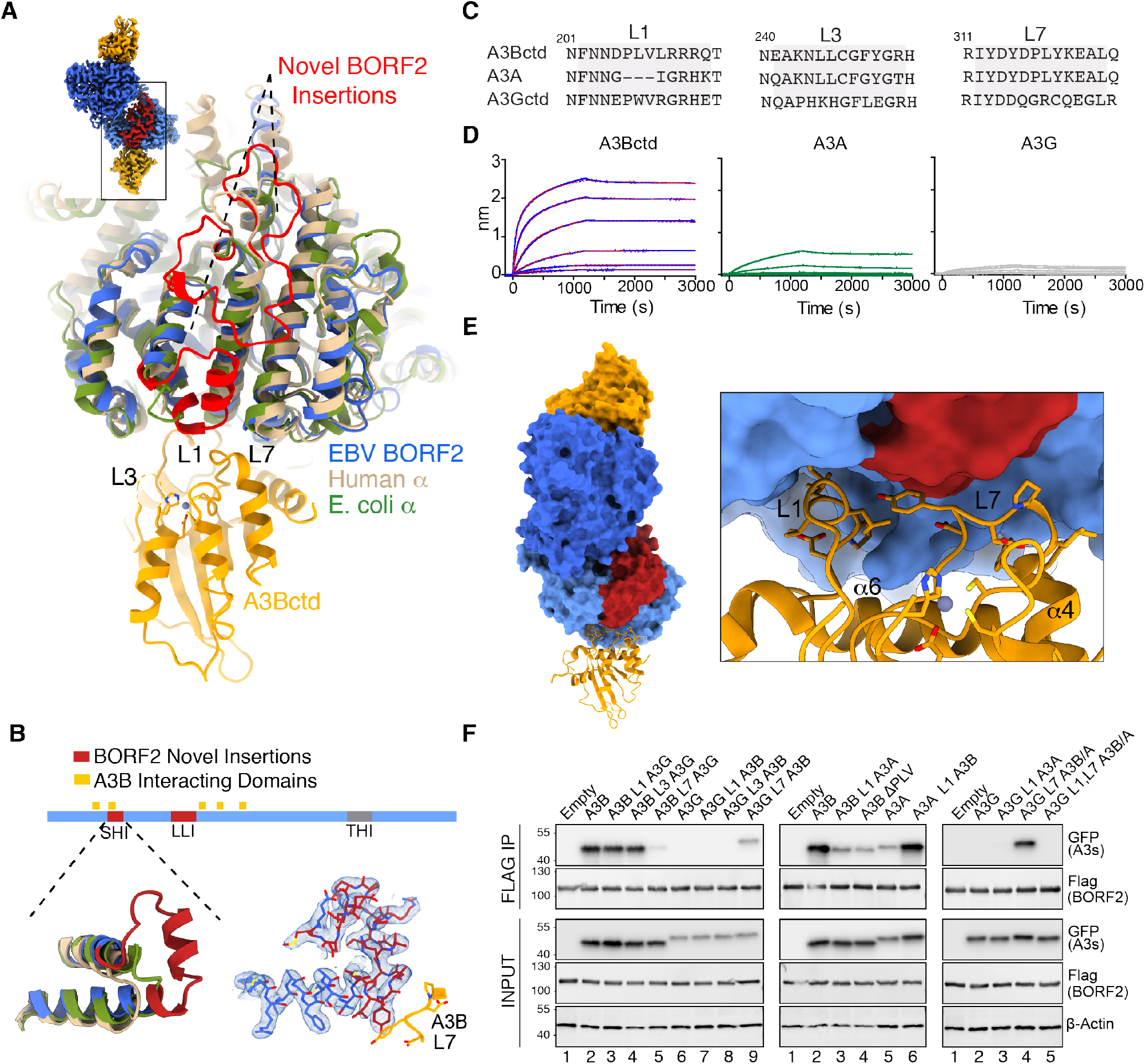
Unique features of EBV BORF2 and its interaction with A3B. **(A)** Overlay of the structures of BORF2 (blue), human RNR α subunit (pdb: 6aui, chain A; tan), and *E. coli* RNR α subunit (pdb:6w4x, chain B; green). Novel BORF2 insertions SHI and LLI are depicted in red. Loop regions surrounding the A3Bctd active site are labeled L1, L3, and L7. **(B)** Schematic of BORF2 (1-826 amino acids) showing the novel SHI and LLI insertions (red) and a triple helix insert (THI) shared with eukaryotic class1a RNRs. Bottom left: a structural comparison of the BORF2 SHI and the corresponding regions of human and *E. coli* RNR α subunits. Bottom right: Cryo-EM map of the BORF2 SHI domain in blue. **(C)** Amino acid alignment of the L1, L3, and L7 regions of A3Bctd, A3A, and A3Gctd (full alignments in fig. S5). **(D)** BLI sensorgrams of BORF2 binding to A3Bctd-mychis, A3A-mychis, and A3G-mychis immobilized on Ni-NTA probes. BORF2-A3Bctd interaction has a calculated K_d_ value of 0.78 based on fitting data with a 1:2 binding model (red-dashed line represents fit of the curves to model). **(E)** A surface representation of the BORF2-A3Bctd complex with one of the A3B protomers represented in ribbons. Right: close-up of the interacting surfaces highlighting BORF2 SHI (red) and A3Bctd L1 and L7 (gold). **(F)** BORF2 (anti-Flag) co-IP experiments with A3Bctd-eGFP, A3A-eGFP, and A3Gctd-eGFP proteins in comparison to the indicated loop swap variants. Input blots are shown below including anti-β-actin as a control.

However, the presence of two novel insertions in EBV BORF2 distinguishes this viral enzyme from all other class Ia RNRs (red in **Fig. 2A-B** and fig. S4). First, a single helix insertion (SHI) is part of an extended loop that packs between four larger structurally conserved helices in BORF2. This novel insertion directly contributes residues to binding A3B (**Fig. 2A-B** and below). Second, a long-loop insertion (LLI) is anchored on the RNR core and positioned to provide stabilizing interactions for the SHI and surrounding helices (red in **Fig. 2A** and fig. S4A-C). In particular, this LLI directly contacts the SHI, which in turn binds to A3B (Fig. S4B-C). These insertions, unique to EBV BORF2, suggest that an evolutionary gain-of-function may have been required to neutralize A3B DNA deaminase activity and create an environment permissive for lytic DNA replication.

All A3s share a common cytidine deaminase fold in which the active site is comprised of a zinc-coordinating domain surrounded by three loops (L1, L3, and L7) that engage substrate single-stranded DNA and can vary significantly between family members [reviewed by (*21*); **Fig. 2A, Fig. 2C**, fig. S5]. There are two key interaction points between BORF2 and A3B. One involving A3B residues 314-320 spanning the L7 region and the other involving residues 203-214 including the L1 region. The BORF2-A3B interaction has a measured dissociation constant of ∼1 nM as determined by BLI experiments (**Fig. 2D**). In comparison, at the same protein concentrations BORF2 has reduced binding for A3A (a near-identical A3 that shares 90% amino acid sequence identity with A3Bctd) and negligible binding for A3G (an A3 with a catalytic domain sharing 64% amino acid sequence identity). The largest differences between A3Bctd, A3A, and A3Gctd are found in the loop regions adjacent to the active site (**Fig. 2A, Fig. 2C**, fig. S5). Indeed, the cryo-EM structure shows that two loop regions of A3Bctd, L1 and L7, have multiple contact points with BORF2 (**Fig. 2E**, fig. S6).

To assess the structural importance of these loops, chimeric A3 proteins with reciprocal loop swaps were tested for binding in co-immunoprecipitation experiments. Replacing L7 of A3B with L7 of A3G nearly abolished A3Bctd pull-down by BORF2 (**Fig. 2F**, left panel, lanes 2 vs 5). Correspondingly, replacing L7 of A3G with L7 of A3B enabled A3Gctd to be pulled-down effectively (**Fig. 2F**, left panel, lanes 6 vs 9). In contrast, replacing L1 or L3 of A3B with the analogous loop regions from A3G or the reciprocal experiment did not seem to have an affect (**Fig. 2F**, left panel). These co-IP experiments demonstrate that L7, as resolved in the structure, is essential for the interaction with BORF2.

A3Bctd and A3A have identical L7 regions, yet BORF2 binds A3Bctd with higher affinity (**Fig. 2A, C-D**). This difference is explained by an important secondary interaction site involving A3Bctd L1. When L1 of A3A is grafted into A3Bctd, or when amino acids 206-PLV-208 are deleted from A3Bctd L1 to make it more A3A-like, marked decreases in A3Bctd co-immunoprecipitation are observed (**Fig. 2F**, middle panel, lanes 2-4). Reciprocally, when L1 of A3Bctd is used to replace L1 of A3A, a clear increase in pull-down strength becomes evident (**Fig. 2F**, middle panel, lanes 5-6). Interestingly, the strong interaction between BORF2 and A3Gctd containing L7 of A3Bctd/A3A is abrogated by replacing the L1 region of this chimera with L1 of A3A (**Fig. 2F**, right panel, lanes 4-5). This can be explained by similarities between A3B and A3G L1 regions where these enzymes both have a 3-residue insertion that is absent in A3A (PLV or PWV, respectively; **Fig. 2C**, fig. S5). These co-IP results combine to explain the importance of these two loop regions and show that BORF2 binds preferentially to A3B.

The BORF2-A3Bctd interface spans a surface area of ∼1040 Å^2^ and involves residues from multiple noncontiguous regions of each protein (**Fig. 2A-B**). The high-affinity interaction between BORF2 and A3B can be explained by the overall nature of the interacting surfaces including multiple hydrophobic and electrostatic contacts (**Fig. 2E**, fig. S6-7). First, two BORF2 tyrosine residues are positioned to interact with residues in the L1 and L7 region of A3B (fig. S7A). Specifically, BORF2 residues Y134 and L133 (SHI domain) directly interact with Y315 in L7 of A3B (fig. S7B). Second, BORF2 residue Y481 is inserted in a hydrophobic pocket generated between the PLV region of A3B L1 and H365 and L369 of the alpha-6 region (fig. S6, S7C). The interlocked positioning of BORF2 Y481 is likely aided by a neighboring proline (P479) that effectively kinks the BORF2 loop containing these residues (fig. S7C). BORF2 residue R484 is situated halfway between residues Y134 and Y481 and provides an electrostatic interaction with A3B residue D314 in L7 and also interacts with BORF2 Y134. Single amino-acid substitutions at these positions in BORF2 or A3B substantially diminish the co-IP interaction (fig. S7D-E).

The n-terminus of human, *E. coli*, and several class 1a RNRs has an evolutionarily conserved mobile ATP cone domain that regulates activity (ATP activates whereas dATP inhibits) [reviewed by (*1*); fig. S8]. The majority of this cone domain is absent in BORF2 including the ATP binding residues and, instead, the n-terminus adopts a helix that mediates formation of a non-canonical and novel dimer (**Fig. 3A-B**, fig. S8A). This dimer is driven by a leucine rich hydrophobic interaction involving two alpha helices from each monomer (*i*.*e*., a four-helix bundle) and electrostatic interactions including residues R39, D26, and E8 (**Fig. 3A**). R39 from each monomer directly interacts with E8 from the same monomer and D26 from the opposing monomer. To investigate whether dimer formation occurs in cells, we co-expressed BORF2 harboring two different tags (n-terminal MBP and c-terminal 3x-Flag) and performed anti-Flag co-immunoprecipitation assays, which resulted in a pulldown of a complex containing both tagged forms of BORF2 (**Fig. 3B**). As predicted by the structure, single amino acid substitutions of R39E or D26R weaken the interaction and, importantly, this defect could be reversed by combining these changes into a double mutant (R39E, D26R) with a restored salt-bridge (*e*.*g*., compare lanes 4-7 in **Fig. 3B**).

**Fig. 3.**
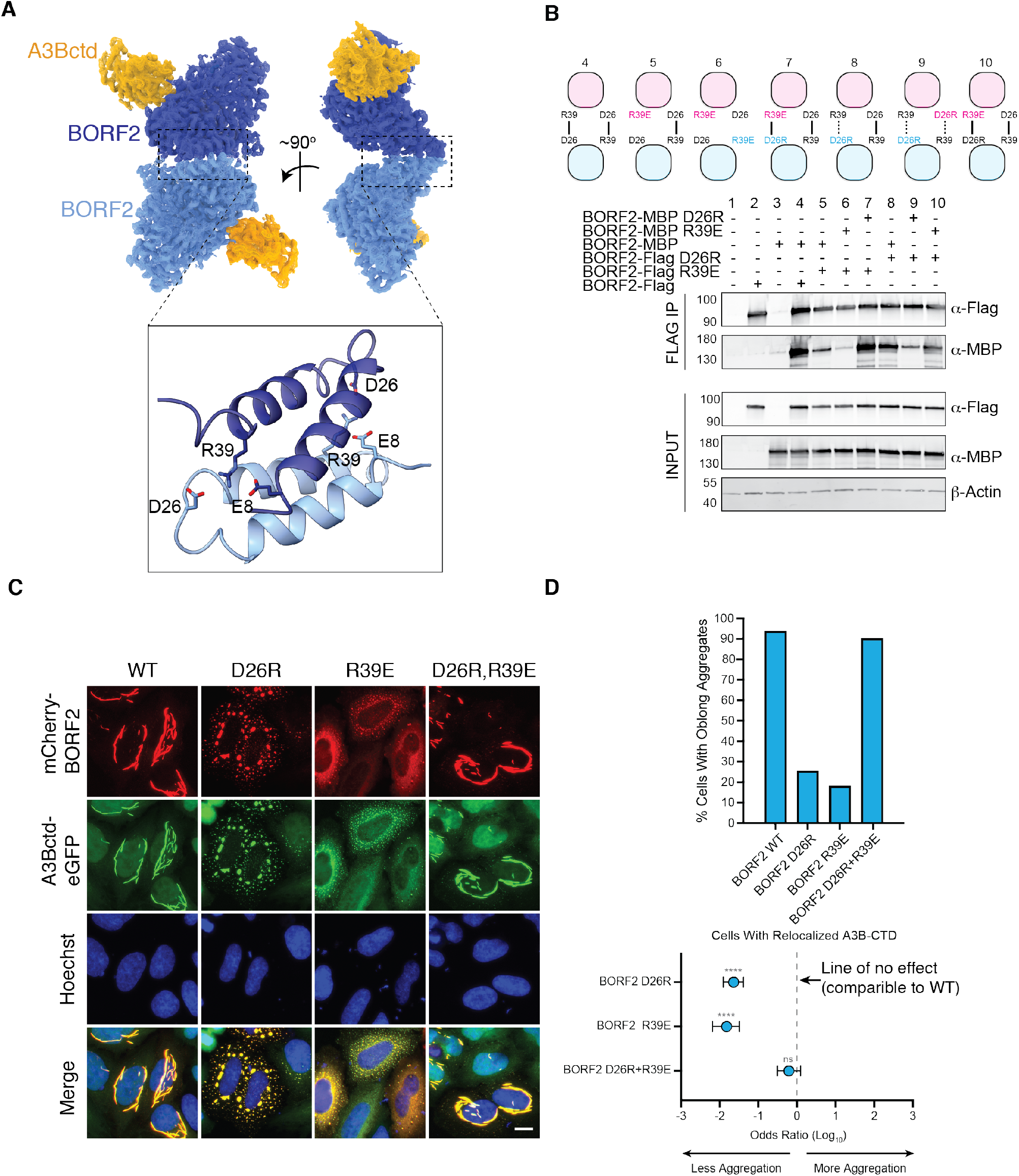
Novel EBV BORF2 dimerization interface is divergent from other RNRs and influences A3B localization pattern. **(A)** Side and front views of the BORF2-A3Bctd structure with a zoom-in of the dimer interface. N-terminal regions of BORF2 form a dimer mediated in part by R39 and E8 of one monomer and D26 of the opposing monomer. **(B)** Co-IP reactions with the indicated BORF2-Flag and MBP-BORF2 constructs. Input blots are shown below including anti-β-actin as a control. Schematics above depict key interactions between BORF2 monomers. **(C)** Representative fluorescent microscopy images of the indicated mCherry-BORF2 and A3Bctd-eGFP constructs (scale = 10µm). Nuclei are stained with Hoechst. **(D)** Quantification of cells forming oblong aggregates upon co-expression of A3Bctd-eGFP and different mCherry-BORF2 constructs (n>100 for each condition). Odds ratios and p-values calculated by Fisher’s exact test (ns, not significant; **** P ≤0.0001). Error bars = 95% confidence intervals.

A feature of the BORF2-A3B interaction is that it results in relocalization of normally nuclear A3B into cytoplasmic aggregates (*8, 9*). We therefore tested to see if BORF2 dimerization influences this localization pattern. Pre-engineered HeLa T-REx cells were transfected with mCherry-BORF2, mCherry-BORF2 R39E, mCherry-BORF2 D26R, and mCherry BORF2 harboring both D26R and R39E, treated 24 hrs later with doxycycline to induce eGFP, A3Bctd-eGFP, or full-length A3B-eGFP, and analyzed by IF microscopy after an additional 24 hrs incubation. Wild-type BORF2 relocalizes both A3Bctd and full-length A3B from the nuclear compartment to the cytoplasm where filament-like aggregates accumulate (**Fig. 3C-D**, fig. S9). BORF2 D26R depletes A3B from the nucleus but does not form similar structures. Likewise, R39E retains the ability to relocalize A3B from the nucleus but has a reduced ability to form these structures. However, the D26R, R39E double substitution restores BORF2’s ability to form these filament-like structures further confirming a direct interaction between these residues. These data indicate that BORF2 dimerization facilitates formation of higher order BORF2-A3B oligomers in cells.

The canonical dimer formed between large subunits of class Ia RNRs is conserved and required for ribonucleotide reduction [(*1, 4*); *e*.*g*., schematics of *E. coli* and human complexes in fig. S8B-C and with yeast RNR in fig. S10A]. EBV carries genes for both the large RNR subunit, BORF2, and the small subunit, BaRF1, and both are expressed during the lytic stage of virus replication (*22-24*). A recent cryo-EM structure of an *E. coli* RNR complex trapped in an active state provided unprecedented detail for RNR catalytic mechanism that involves orchestrated radical transfer between the small subunit (β) and large subunit of the RNR (α) (*25*) (fig. S3B, S8B). A structural overlay of this *E. coli* RNR α_2_β_2_ complex and a monomer of the BORF2-A3Bctd complex shows that the A3B binding site is on the opposite side and is unlikely to overlap with the predicted BaRF1 interaction domain (fig. S10B). Furthermore, considering that the BORF2 predicted catalytic residues are highly conserved with the analogous *E. coli* RNR residues (fig. S3), it is likely that the BORF2-BaRF1 interaction would be positioned in a similar fashion to the *E. coli* large and small subunits, respectively (fig. S10B). Ribonucleotide reductase activity may therefore require the formation of a canonical dimer between BORF2 subunits as depicted in Fig. S10C-D (in contrast to the novel n-terminal region-mediated dimer observed here in the cryo-EM structure). This predicted canonical dimerization region is disordered in the BORF2-A3Bctd structure described here (fig. S10C). Given the flexibility detected in the complex (movies S1 and S2) and that large subunits of RNRs show dynamic oligomerization profiles depending on the active state of the RNR [reviewed by (*4*)], it is possible that this region may become better ordered upon canonical dimer formation.

Here, we report a cryo-EM structure of the large RNR subunit of EBV, BORF2, in complex with the A3B catalytic domain. This provides the first structural explanation for a non-ribonucleotide reducing role for RNRs in biology and additional insights into oligomerization permutations exhibited by these large subunits that have relevance to multiple functions. The structure reveals the overall architecture of the viral RNR and how it has evolved a unique mechanism to selectively bind and inhibit A3B as opposed to other highly homologous family members. Even A3A, which has the highest overall identity with the catalytic domain of A3B (>90%), is a less preferred substrate due to differences in loop 1 (**Fig. 2**). This is in contrast to HIV-1 Vif counteraction mechanism in which Vif binds to and facilitates the degradation of up to five different APOBEC3s (*11*). The striking specificity of BORF2 for A3B may be explained by A3B expression levels in B lymphocytes (*26*), which are a primary cell type for EBV pathogenesis (*27*). The atomic details of the BORF2-A3B interaction may therefore inform the development of antivirals that block this interaction and allow A3B to restrict the replication of EBV and related herpesviruses. In addition, given the prominent pathological role of A3B in cancer mutagenesis (*12, 13*) insights from the potent A3B inhibitory mechanism of BORF2 could lead to strategies to slow or stop the evolution of A3B-positive tumors and improve clinical outcomes.

## Acknowledgments

We would like to thank Hideki Aihara for providing constructs, Wei Zhang and Momoko Shiozaki for grid preparation and screening assistance, Xiaowei Zhao for help with initial dataset collection, Margaret Brown for technical assistance in early stages of the project, William Brown for providing expertise with BLI experiments, and Matthew Jarvis for computational assistance.

## Author contributions

NMS & RSH initiated the project and lead the studies. NMS conceptualized and designed the studies with input from authors. NMS purified proteins and performed biochemical and BLI assays. NMS and SNM generated constructs. NMS & RY prepared and screened grids. RY collected cryo-EM data. ZY supervised cryo-EM data collection. JSM, RY, NMS & ZY processed cryo-EM datasets. JSM processed the final cryo-EM data and calculated the deposited maps. NMS and KS built and refined the structural model with input from JSM. SNM, AZC & NMS performed immunoprecipitation assays. SNM generated cell lines and performed and developed methods to quantify localization experiments. MAC provided cell culture and technical expertise. NMS & RSH wrote the manuscript with contributions from all authors.

## Funding

This work was supported in part by NCI P01-CA234228 (to RSH), NIAID R56-AI150402 (to NMS), and by the University of Minnesota Masonic Cancer Center, Academic Health Center, and College of Biological Sciences. Contributions of JSM supported by Welch Foundation grant number F-0003-19620604. Salary support for SNM was provided by NIAID T32-AI083196 and subsequently NIAID F31-AI161910. RSH is the Margaret Harvey Schering Land Grant Chair for Cancer Research, a Distinguished University McKnight Professor, and an Investigator of the Howard Hughes Medical Institute.

## Competing interests

The authors declare no competing interests.

## Data and materials availability

Cryo-EM maps and a structural model have been deposited in the EMDB and PDB with accession codes EMD-24716, EMD-24715, EMD-24709, and pdb:7wr6, respectively. All other data and materials are available upon request.

## Supplementary Materials

Materials and Methods

Figs. S1 to S10

Table S1

Movies S1 to S2

References (*1*–*27*)

## Notes

### Competing Interest Statement

The authors have declared no competing interest.

