## Supplementary Methods, Table S1, and Figs S1-S10 for "CryoEM structure of the EBV ribonucleotide reductase BORF2 and mechanism of APOBEC3B inhibition"

**This PDF file includes:**

Materials and Methods
Figs. S1 to S10
Table S1
References
Captions for Movies S1 to S2

**Other Supplementary Materials for this manuscript include the following:**

Movies S1 to S2

### Materials and Methods

#### Bacterial and human cell expression constructs

*E. coli* expression vectors with an n-terminal MBP tag or 6x-HisSumo tag and a codon-optimized BORF2 gene were gifts from Hideki Aihara. To create the MBP-BORF2 (1-826) human cell expression vector, the MBP-codon optimized BORF2 was PCR amplified from the bacterial expression constructs above using primers 5’ AAG CTT GGT ACC ACC ATG AAA ATC GAA GAA GGT AAA CTG GTA ATC TGG ATT AAC G and 5’ CTA GAC TCG AGT CAC TGG CAG CTT TCG CAC GCG CGT TCA G and ligated into pcDNA3.1 (Invitrogen) using restriction enzymes KpnI-HF (NEB #R3142) and XhoI (NEB #R0146). This construct was used to express protein for determining the cryo-EM structure, BLI experiments in **Fig. 2**, and for co-IP experiments described in **Fig. 3B**. The MBP-BORF2 R39E, D26R, and R39E, D26R substitutions were made by site-directed mutagenesis. These constructs were used for co-IP experiments described in **Fig. 3B**.

The n-terminal tagged mCherry and c-terminal 3x flag tagged BORF2 used in localization and co-IP studies were cloned into pCDNA5TO and pCDNA4TO (Invitrogen). The BORF2 starting methionine in the mCherry-BORF2 construct is deleted to prevent internal translation initiation. All mCherry-BORF2 and BORF2-3xflag single-point amino acid substitutions were generated by site-directed mutagenesis.

The A3-mychis constructs used to express protein for structural work, biochemical assays, and BLI assays have been described (*28*). Briefly, A3Bctd (193-382)-mychis, A3A (1-199)-mychis, and A3G (1-384)-mychis constructs are in pcDNA3.1 c-terminal mychis vectors (Invitrogen). The A3A and A3B coding sequence is disrupted by an intron to avoid toxicity in *E. coli* during cloning and plasmid preparation.

Full-length A3B-eGFP, A3Bctd$_{193-382}$-eGFP, full-length A3A-eGFP, A3Bctd$_{193-382}$-eGFP used in localization experiments (**Fig. 3C**, fig. S9) and co-IP experiments (**Fig. 2E**, fig. S7) were cloned into pCDNA5TO (Invitrogen) using PCR amplification, restriction enzyme digestion, and ligation. The loop swap constructs A3Bctd$_{193-382}$A3AL1$_{25-30}$, A3Bctd$_{193-382}$L1$_{deltaPLV}$, A3Bctd$_{193-382}$A3GL1$_{209-217}$, A3A_A3BL1$_{205-213}$, A3A_A3GL1$_{209-213}$, and A3A_A3GL3$_{247-254}$ were made by site-directed mutagenesis.

Unless otherwise stated, Pfu-Ultra II was used for site-directed mutagenesis and HF-Phusion (M0530) for all other PCR reactions. All constructs were sequence confirmed using Sanger sequencing. Wild-type constructs match the following Genbank accessions: BORF2 (V01555.2), A3B (NM_004900), A3A (NM_145699), and A3G (NM_021822).

#### Protein expression and purification

Full-length BORF2 was expressed in 293T cells as an n-terminal maltose binding protein (MBP) tagged fusion. 293T cells were grown at 37ºC $CO_2$ in RPMI media supplemented with 10% fetal bovine serum and 1X antibiotic-antimycotic solution (Gibco Life Technologies).

Between 10-30 15cm$^3$ plates of 293Ts were transfected with 10-20 µgs of plasmid using a 3:1 ratio of PEI to plasmid DNA. The next day, fresh media was exchanged and the cells were incubated for an additional 36-48 hrs. Cells were harvested and frozen in -80ºC and thawed and resuspended in lysis buffer (50mM Tris-HCl pH 7.4, 300 mM NaCl, 10% glycerol, 1 tablet EDTA-free protease inhibitor, and 20 µg/ml RNase A). The resuspension was sonicated on an ice bath 2-3 times using a Branson sonifier (Duty cycle: 4, output 4, 2min pulses). Cellular debris was removed by centrifugation at 15,000xg for 45 min at 12ºC. MBP-BORF2 was purified by binding to amylose resin (NEB). Resin was washed with 50 mM Tris-HCl, 300 mM NaCl, 10% glycerol, and MBP-BORF2 was eluted from the resin with 100 mM maltose. TCEP (final concentration 1mM) was added to the elution and the protein was concentrated to ~0.5 mls. The sample was injected into a S200 Increase 30/300 (GE Healthcare) preequilibrated with 20 mM Tris HCl pH 8 and 150 mM NaCl. Purified complex corresponding to the peak fraction was collected, TCEP was added (1 mM), and the protein was concentrated prior to experimentation.

A3Bctd-mychis and A3A-mychis proteins were expressed by transfecting 10-30 15cm$^3$ plates of 293Ts with up to 10 µg of plasmid using a ratio of 3:1 PEI to plasmid DNA. The cells were harvested ~48 hrs post-transfection. Pellets were frozen in -80ºC. The pellets were thawed and resuspended in 50 mM Tris HCl pH 7.4/8, 500 mM NaCl, 10% glycerol, 5 mM imidazole, 1 tablet EDTA-free protease inhibitor, and 20 µg/ml RNase A. Cells were lysed in an ice-bath by sonication using a Branson sonifier (Duty cycle: 4, output 4, 2 min pulses, 2-3 times) followed by centrifugation (15,000xg, 45min, 12ºC). Supernatants were applied to Talon metal affinity resin (Clontech Takara) to capture the his-tagged proteins. The resin was washed with 50 mM Tris HCl pH 8, 500 mM NaCl, and 5mM imidazole. Bound proteins were eluted with 50 mM Tris HCl pH 7.4/8, 500 mM NaCl, and 250mM imidazole. TCEP was added to the elution (final concentration 1 mM) and the sample was concentrated to 0.5 mls. The sample was further purified by size-exclusion chromatography using an S200 Increase 30/300 (GE Healthcare/Cytivia) that was preequilibrated with 20mM Tris HCl pH 8 and 300 mM NaCl. A3G-mychis was purified similarly except with an additional RNase A treatment prior to SEC.

The MBP-BORF2/A3B-mychis complex used to determine the cryo-EM structure was purified by mixing excess purified A3B-mychis with MBP-BORF2 (after amylose resin purification). The complex was isolated by size-exclusion chromatography in buffer containing 20 mM Tris-HCl pH 8 and 150 mM NaCl. TCEP was added to a final concentration of 1 mM and the complex was concentrated and stored at -80ºC.

**DNA deaminase assays**

Purified A3Bctd-mychis was diluted to 100 nM in reaction buffer containing 10 mM Hepes pH 7.4, 5 mM EDTA. 50 mM NaCl, and 0.5 mM TCEP. MBP-BORF2 was serially diluted in the same buffer to concentrations of 200, 100, and 50 nM. Equal volumes of MBP-BORF2 and A3Bctd-mychis were mixed together prior to adding oligonucleotide. The oligonucleotide with a 3' FAM used as a substrate for A3Bctd was ordered from IDT and contains the preferred A3B trinucleotide sequence "TCA" (ATT ATT ATT ATT CAA ATG GAT TTA TTT ATT TAT TTA TTT ATT T-FAM). The oligonucleotide was diluted to 800 nM in reaction buffer. Equal volumes of substrate and protein complex were mixed together and incubated for 1 hr at 37 ºC (total reaction volume 10 µl). This was followed by addition of UDG (NEB M0280L) for 10 min. to remove the uracil base resulting from A3B deaminase activity. The reaction mix was then incubated with 1M NaOH for 10 min at 98ºC to break the phosphodiester backbone at the abasic site. 11 ul of loading dye was added to each reaction, and reaction products were separated on a 15% TBE urea acrylamide gel. The gel was imaged using a Typhoon7000 (GE Healthcare).

**Biolayer interferometry (BLI) assays**

BLI assays were performed using an Octet RED (FortéBio). A3Bctd mychis, A3A mychis, and full-length A3G mychis were loaded onto Ni-NTA biosensors (FortéBio cat #185101) at 12.5 µg/ml for A3Bctd and A3A, and 8 µg/ml for A3G. MBP-BORF2 was serially diluted (200, 100, 50, 25, 12.5, and 6.25 nM). All proteins were diluted in buffer containing 20 mM Tris-HCl pH 8 and 300 mM NaCl (reaction buffer). The assay was performed in 384 tilted bottom microplates (FortéBio Cat #18-5076). 60 µl volumes of reaction buffer or protein samples were used in each condition. Prior to running the assay, the biosensor probes were hydrated in reaction buffer for at least 10 min. The assay was set up in the following order 1) equilibration in buffer, 2) load A3-mychis proteins (300 sec), 3) wash/equilibrate in buffer, 4) association with BORF2 (1200 sec), and 5) dissociation in buffer (1800 sec). The reaction temperature was set at 30ºC. The results were analyzed using ForteBio Octet data analysis software 11.1.2. Wells containing no BORF2 were assigned as the reference sample. The data best fit a 1:2 binding model.

**Cell line constructions**

HeLa cells were transduced with a pLENTI6 TetRepressor construct and selected with 5 µg/ml blasticidin (GoldBio #B-800-25). The cells were subcloned by serial dilution to obtain single cell clones. The clonal populations were stably transfected with pcDNA5TO vectors containing eGFP, A3Bctd-eGFP, or A3B-eGFP. The cells were selected with hygromycin (GoldBio #H-270-1) and subcloned by serial dilution. Representative clones were used for IF experiments (**Fig. 3C**, fig. S9).

**Immunofluorescence microscopy experiments**

HeLa T-REx cells engineered to stably express eGFP, A3Bctd-eGFP, and A3B-eGFP were grown in RPMI media and used for IF experiments described in **Fig. 3C** and fig. S9. Approximately, 3000 cells per well were plated in 96-well imaging plate (Coning #3904). The next day cells were transfected with 25 ng pcDNA5TO-mCherry-BORF2, pcDNA5TO-mCherry-BORF2 D26R, pcDNA5TO-mCherry-BORF2 R39E, or pcDNA5TO-mCherry-BORF2 D26R, R39E constructs using Trans-IT LTI (Mirus #MIR 2300). The cells were treated with 50 ng/µl doxycycline approximately 24 hrs later. The following day, cells were washed with 1X PBS, fixed using 4% formaldehyde (ThermoFisher #28906) and the nuclei were stained with Hoechst 33342 (ThemoFisher #62249). Images were collected using a Cytation1 (BioTek). A total of 50 images per well were collected at 20x magnification using the automated imaging function.

Background subtraction was performed using Fiji (*29*) and images were analyzed using CellProfiler v4.2 (*30*). A pipeline was generated to trace cells, segment the cytoplasmic and nuclear compartments, and identify aggregates/oblong structures in the cytoplasm. Images showing cell segmentation of nuclear and cytoplasmic compartments (middle; green and yellow tracing) and aggregate and oblong structures (right; pink tracing) are shown below. At least 100 cells were quantified for each condition. CellProfiler Analyst v2.2.1 (*31*) was used for scoring phenotypes by machine learning in order to quantify the population of cells within each condition with relocalized A3Bctd-eGFP, full-length A3B-eGFP, eGFP, and/or aggregates/oblong structures (**Fig. 3C**, fig. S9).

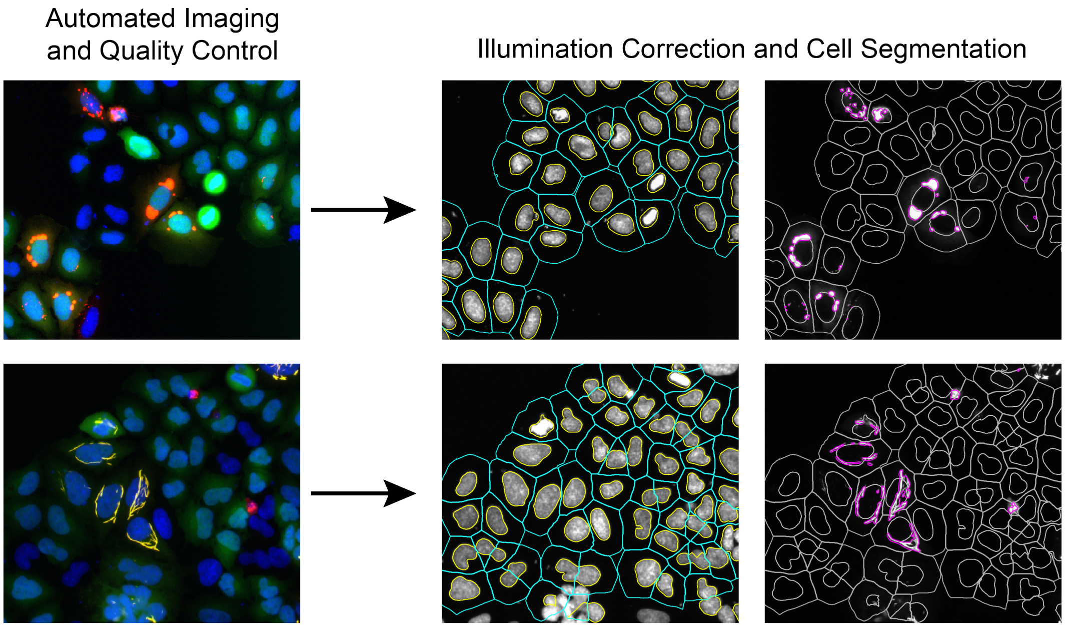


### Immunoprecipitation assays

Immunoprecipitation assays descripted in **Fig. 2E**, **Fig. 3B**, and fig. S7 were performed as described (*8*) with a few adjustments. Briefly, 250,000 293T cells in 6 well plates were transfected with 100 ng of each construct using TransIT-LTI (Mirus #MIR 2304). Cells were harvested ~48 hrs after transfection and resuspended in 700 µl lysis buffer (50 mM Tris-HCl pH 7.5, 150 mM NaCl, 10% glycerol, 0.5% tergitol, 1 tablet EDTA-free protease inhibitor tablet, and 100 µg/ml RNase A). Resuspension was sonicated on ice-bath for 10 sec at the lowest setting using a Branson sonifier and then centrifuged at 15,000 rpm for 10 min. A 30 µl aliquot of supernatant was removed for input detection. Remaining supernatant was incubated with 25 µl of resin anti-FLAG magnetic beads (Sigma #M8823) per sample overnight with gentle rotation at 4ºC. Samples were washed 3 times with lysis buffer and eluted with 30 µl of elution buffer (50 mM Tris-HCl pH 7.5, 150 mM NaCl, 0.05% tergitol, 10% glycerol, flag peptide). The immunoprecipitation assays described in **Fig. 3B** were done following the same protocol with the exception that 400,000 293T cells were transfected with 200 ng of each listed construct.

Elution and input for assays described in **Fig. 2E** were evaluated by immunoblotting with rabbit anti-flag at 1:5,000 to detect BORF2-3xflag (sigma #F7425), mouse anti-GFP at 1:5,000 dilution to detect A3-GFP (Living Colors #JL-8), and rabbit anti-actin at 1:5,000 (Cell Signaling #13E5) to detect actin (loading control). Secondary antibodies used were IRDye 680RD anti-Rabbit (LI-COR #925-68071) and HRP-linked anti-Mouse (Cell Signaling #7076), both at 1:10,000 dilution. Elutions and inputs described in **Fig. 3B** were detected by immunoblot using mouse anti-flag at 1:4000 dilution, rabbit anti-MBP (Invitrogen #PA1.989) at 1:3000 dilution, and rabbit anti-actin listed above at 1:5000 dilution. Secondary antibodies used were IRDye

680RD anti-Rabbit (LI-COR #925-68071) and IRDye 800CW anti-mouse (LI-COR #926-32210) at 1:20,000 dilution.

**Cryo-EM grid preparation**

Quantifoil grids R-1.2/1.3 Cu 400 mesh (Electron Microscopy Sciences) were glow discharged for 1 min at 15 mA using a PELCO easiGlow apparatus. 3 µl of sample at 0.33 mg/ml was applied to the grid and a FEI Vitribot Mark IV was used to plunge-freeze grids with the following parameters (temperature 6ºC, humidity 95%, blotting force 3, blotting time 3). After blotting, the grid was rapidly frozen in liquid ethane.

**Cryo-EM data collection**

A total of 7,488 movies were collected from a frozen grid imaged on a Titan Krios cryo-electron microscope (FEI/Janelia Research Campus) operating at 300keV and equipped with a spherical aberration corrector, an energy filter (Gatan GIF Quantum) and a post-GIF Gatan K3 direct electron detector. Movies were acquired at a calibrated magnification of x59,242 on the K3 camera in CDS mode, corresponding to 0.844 Å per physical pixel (0.422 Å per super-resolution pixel). The dose rate on the specimen was set to be 11.5 electrons per $Å^2$ per second and total exposure time was 5.22 s. 50 frames per movie were collected at 1.2 e-/$Å^2$ dose per frame for a total of 60e-/$Å^2$ dose per movie. The nominal defocus range was set at -0.8 to -2.5 µm. Camera gain reference map was taken at the start of the data collection session but not applied to each movie series to limit file size. To further save disk space, each movie series was saved in tiff format with LZW compression. Data were collected using SerialEM. Data collection parameters are shown in table S1.

**Cryo-EM data processing**

Cryo-EM data were processed in cryoSPARC v3.2.0 (*15*) as depicted in the cryo-EM workflow (fig. S1). After movie alignment and CTF estimation, cryoSPARC blob picker and 2D classifications were performed to select good templates for template-based particle picking. Particles were extracted (3x-fourier crop) followed by several rounds of 2D classifications. The 2D classes showed the presence of monomeric and dimeric species of the complex. The classes corresponding to the dimeric species were selected for Ab Initio reconstruction and heterogeneous refinement. Particles were then re-extracted without fourier cropping followed by additional rounds of 2D classifications to remove suboptimal particles. Good classes were selected for Ab initio reconstruction and heterogeneous refinement resulting in a 2.8 Å reconstruction with C2 symmetry imposed. The particles from this reconstruction were symmetry expanded. A mask around the BORF2-A3B monomer was generated using ChimeraX (*32*), and local refinement was performed with the symmetry expanded particle stack yielding a 2.55 Å map. A composite map from this focused refinement was generated in phenix v1.18-3855-000 [combine focused map feature (*33*)] to represent the dimeric form of the complex. The focused refined and composite map were sharpened using DeepEMhancer (*34*). The MBP tag on the n-terminus of BORF2 was not detected in any of the maps.

**Model building and refinement**

Crystal structure of A3Bctd [pdb 5cqh (*28*)] and a model of BORF2 generated using Phyre2 (*35*) were docked and fitted into the cryo-EM maps using ChimeraX (*32*). The model was further corrected and built manually in coot (*36*) and refined using real space refinement in Phenix (*37*). Model refinement statistics are shown in Table s1. All figures of the model were made using pymol (DeLano Scientific), Chimera (*38*), or ChimeraX (*32*).

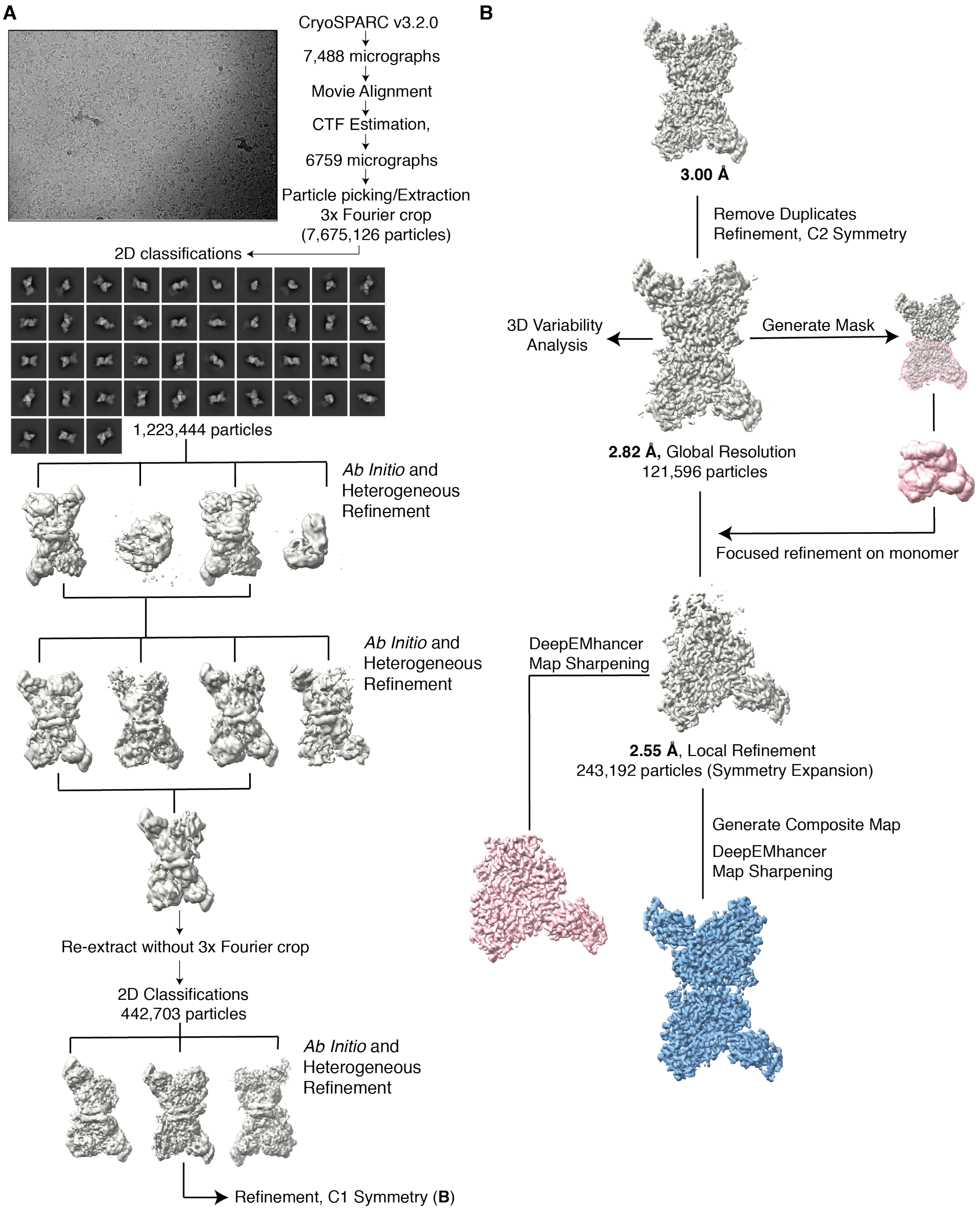
A
CryoSPARC v3.2.0
7,488 micrographs
Movie Alignment
CTF Estimation,
6759 micrographs
Particle picking/Extraction
3x Fourier crop
(7,675,126 particles)
2D classifications
1,223,444 particles
*Ab Initio* and
Heterogeneous
Refinement
*Ab Initio* and
Heterogeneous
Refinement
Re-extract without 3x Fourier crop
2D Classifications
442,703 particles
*Ab Initio* and
Heterogeneous
Refinement
Refinement, C1 Symmetry (**B**)
B
**3.00 Å**
Remove Duplicates
Refinement, C2 Symmetry
3D Variability
Analysis
Generate Mask
**2.82 Å,** Global Resolution
121,596 particles
Focused refinement on monomer
DeepEMhancer
Map Sharpening
**2.55 Å**, Local Refinement
243,192 particles (Symmetry Expansion)
Generate Composite Map
DeepEMhancer
Map Sharpening

**Fig. S1. Cryo-EM data processing work-flow.**
(**A**) Top left: representative micrograph and 2D classes with workflow of *Ab initio* reconstruction and refinement.
(**B**) Refinement workflow for reconstruction selected in panel A.

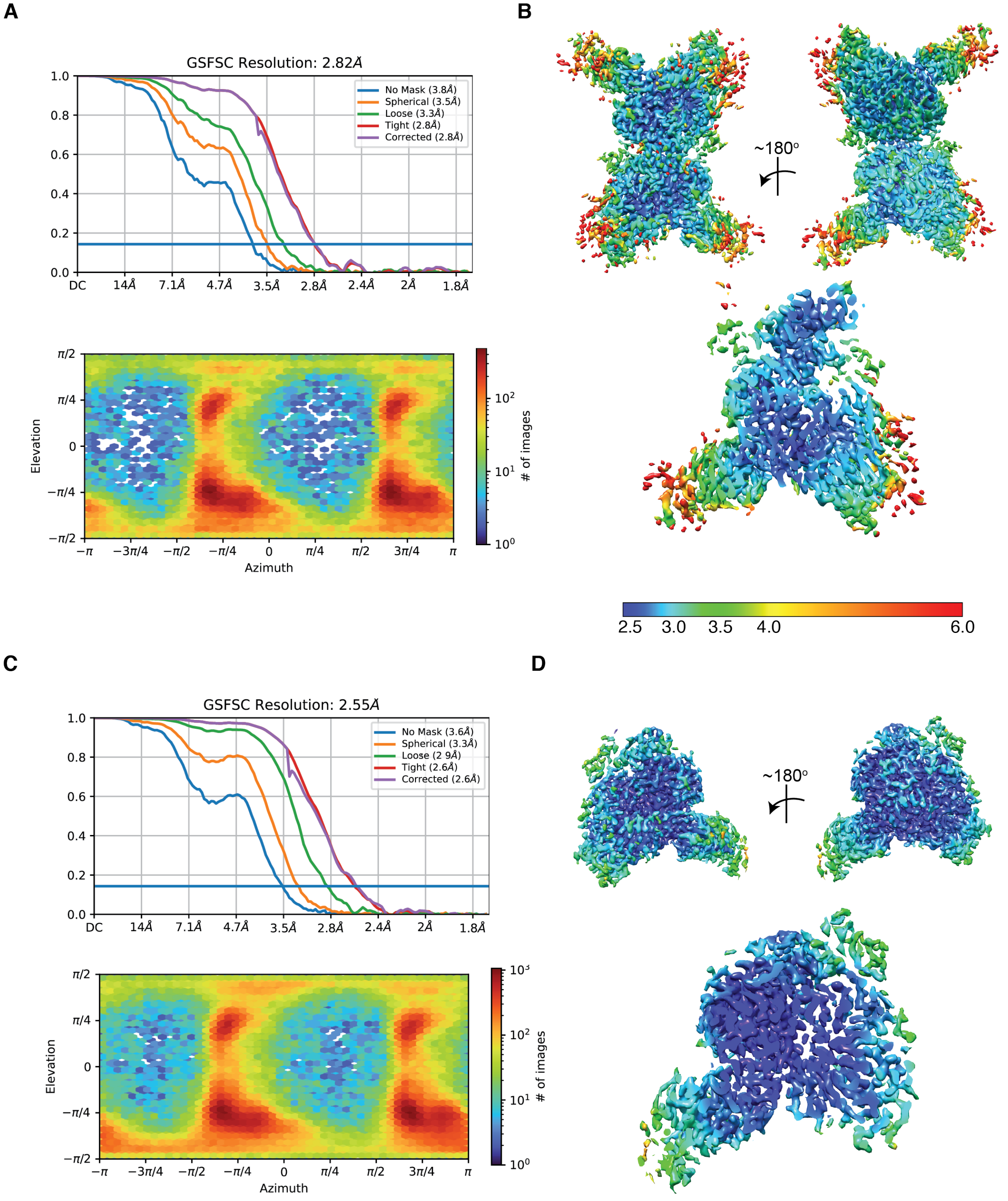


**Fig. S2. Cryo-EM data validation**
(**A**) Top: FSC curves for the 2.8 Å global resolution cryo-EM reconstruction (fig. S1). Bottom: viewing direction distribution plot.
(**B**) Top: Two views of the 2.8 Å global resolution cryo-EM map colored to depict local resolution range. Bottom: close up of the BORF2-A3B monomer and central slice through one of

the views showing the local resolution range. Color key shown on bottom. Resolution is in angstroms. Local resolution calculated using local estimation function in cryoSPARC.
(**C**) Top: FSC curves of 2.55 Å local refinement reconstruction (fig. S1). Bottom: viewing direction distribution plot.
(**D**) Top: Two views of the 2.55 Å local refined cryo-EM map colored to depict local resolution range. Bottom: zoom in and central slice through one of the views showing the local resolution range. Color key shown on the bottom of panel B. Resolution is in angstroms. Local resolution calculated using local estimation feature in cryoSPARC.

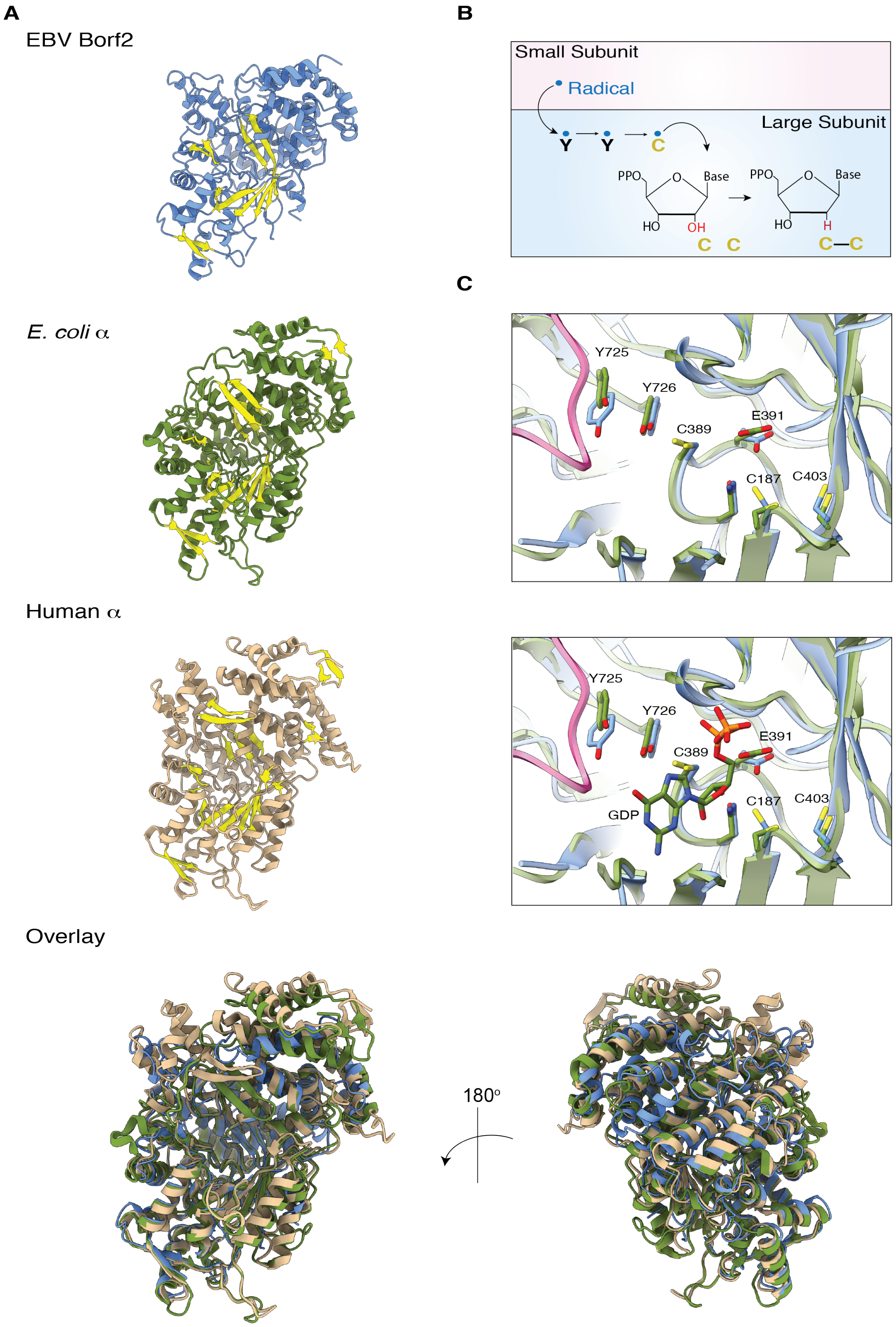

A
EBV Borf2
E. coli α
Human α
Overlay
180°
B
Small Subunit
Radical
Large Subunit
Y
Y
C
PPO
O
Base
HO
OH
C C
PPO
O
Base
HO
H
C—C
C
Y725
Y726
C389
E391
C187
C403
Y725
Y726
E391
C389
C187
C403
GDP

**Fig. S3. RNR large subunit structural conservation.**

(**A**) EBV BORF2 (blue), *E. coli* RNR α-subunit (pdb 6w4x; green), and human RNR α-subunit (pdb 6aui; tan) share a conserved α/β barrel catalytic core and overall architecture. Beta-strands are highlighted in yellow for visual comparison, and overlays are shown below.
(**B**) A cartoon schematic of the proposed radical-based mechanism for ribonucleotide reduction with coordination between the small (pink) and large (blue) subunits [see recent reviews (*1, 5*)].
(**C**) A zoom-in of the overlay of the active site regions of EBV BORF2 and (blue) and *E. coli* RNR α-subunit (pdb 6w4x chain B; green) showing conservation of the tyrosine and cysteine residues involved in radical transfer and oxidation. Numbering of labelled residues corresponds to BORF2.

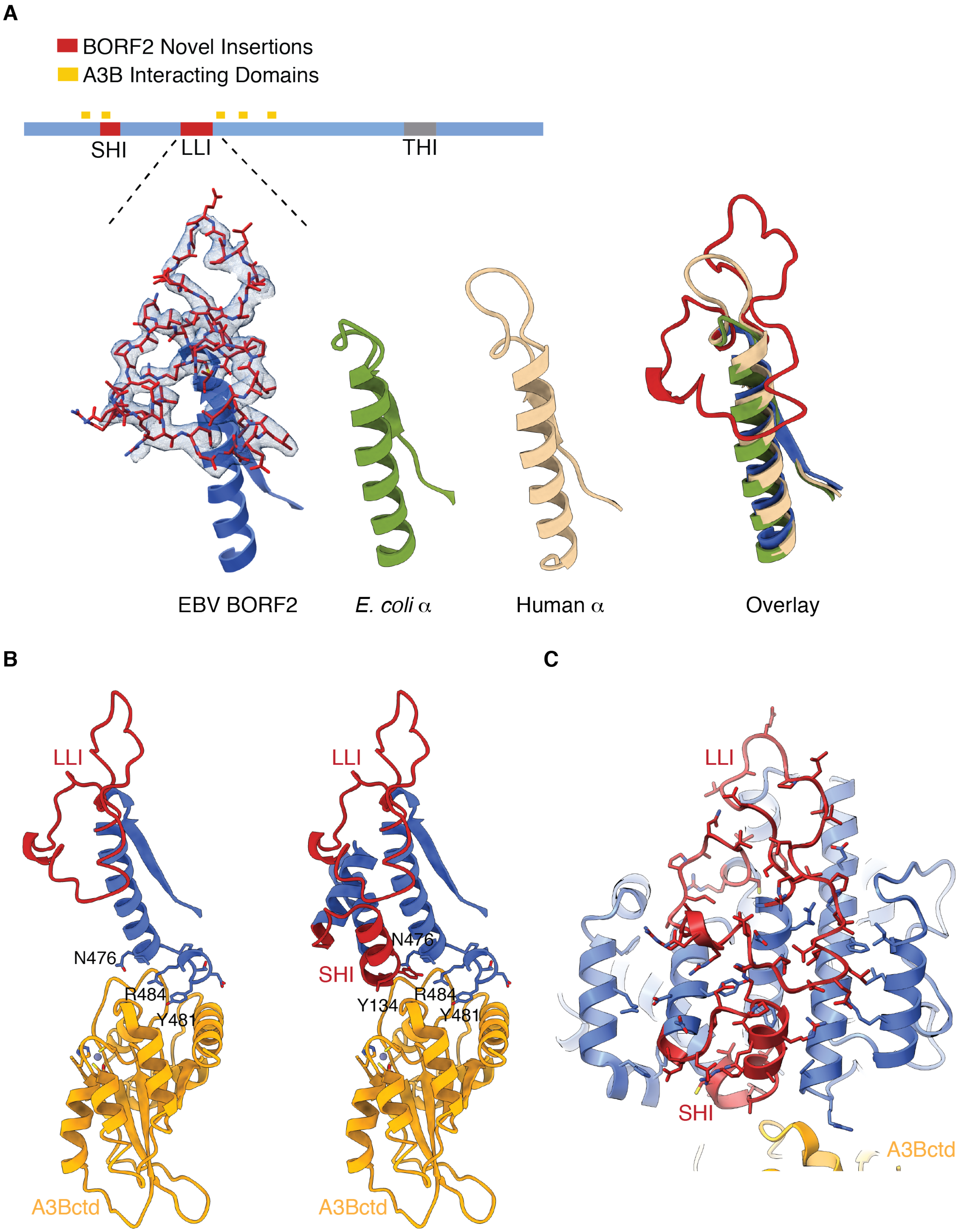
A
BORF2 Novel Insertions
A3B Interacting Domains
SHI
LLI
THI
EBV BORF2
E. coli α
Human α
Overlay
B
LLI
N476
R484
Y481
A3Bctd
LLI
N476
SHI
Y134
R484
Y481
A3Bctd
C
LLI
SHI
A3Bctd

**Fig. S4. Novel insertions in EBV BORF2.**

(**A**) Schematic of BORF2 (1-826 amino acids) showing the novel SHI and LLI insertions (red), and a triple helix insert (THI) shared with eukaryotic class1a RNRs. Select view of the BORF2 LLI insertion and the corresponding regions of the human RNR α subunit (pdb: 6aui, chain A; tan) and the *E. coli* RNR α subunit (pdb:6w4x, chain B; green). Cryo-EM map (bottom left) is shown for the LLI region of BORF2 and is represented by blue mesh.

(**B**) Ribbon schematics showing connectivity between BORF2 insertions LLI and SHI and A3Bctd. The left schematic emphasizes core interactions and the right schematic provides additional BORF2 structure including the SHI. BORF2 residues that interact with A3Bctd are labeled.

(**C**) A faded ribbon schematic showing the extensive interactions that occur both between the SHI and LLI and between these novel insertions and the protein core.

**A**

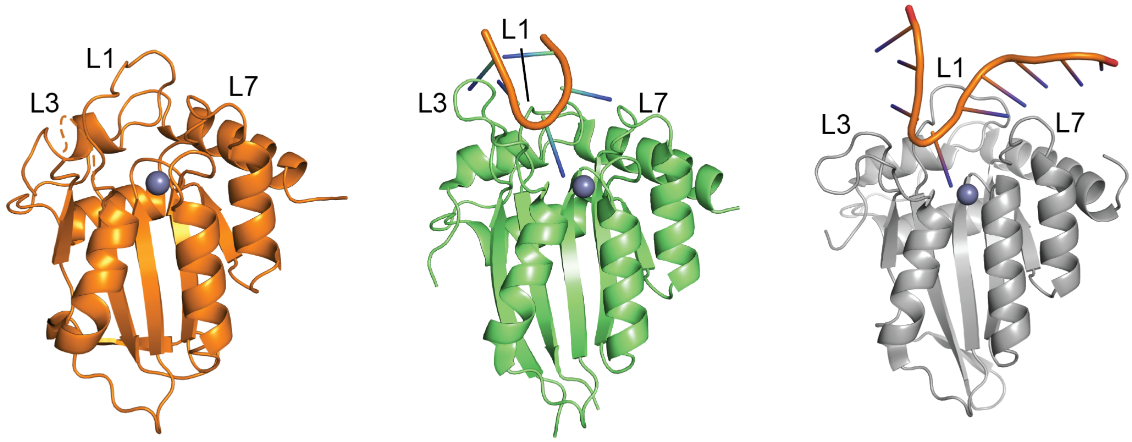


A3Bctd (this study)    A3A-ssDNA (pdb: 5sww)    A3Gctd-ssDNA (pdb: 6bux)

**B**

```
                                      L1
A3Bctd     ---------RYLMDPDTFTFNFNNDPLVLRRRQTYLCYEVERLDNGTWVLMDQHMGFLCN 240
A3A        MEASPASGPRHLMDPHIFTSNFNNG---IGRHKTYLCYEVERLDNGTSVKMDQHRGFLHN 57
A3Gctd     ---------RHSMDPPTFTFNFNNEPWVRGRHETYLCYEVERMHNDTWVLLNQRRGFLCN 240
                    *:  ***   **  ****       *::*********:.*.*  *  ::*:  *** *

                L3
A3B        EAKNLLCGFYGRHAELRFLDLVPSLQLDPAQIYRVTWFISWSPCFSWGCAGEVRAFLQEN 300
A3A        QAKNLLCGFYGRHAELRFLDLVPSLQLDPAQIYRVTWFISWSPCFSWGCAGEVRAFLQEN 117
A3G        QAPHKHGFLEGRHAELCFLDVIPFWKLDLDQDYRVTCFTSWSPCFS--CAQEMAKFISKN 302
           :*  :     :  ****** ***::*   :**   *  ****  *  *******   **  *:   *:.:*

                                   L7
A3B        THVRLRIFAARIYDYDPLYKEALQMLRDAGAQVSIMTYDEFEYCWDTFVYRQGCPFQPWD 360
A3A        THVRLRIFAARIYDYDPLYKEALQMLRDAGAQVSIMTYDEFKHCWDTFVDHQGCPFQPWD 177
A3G        KHVSLCIFTARIYDDQGRCQEGLRTLAEAGAKISIMTYSEFKHCWDTFVDHQGCPFQPWD 362
           .** * **:***** :   :*.*: * :***::*****.**::****** :*********

A3B        GLEEHSQALSGRLRAILQNQGN      382
A3A        GLDEHSQALSGRLRAILQNQGN      199
A3G        GLDEHSQDLSGRLRAILQNQEN      384
           **:**** ************ *
```

**Fig. S5. APOBEC3 structure and protein sequence alignment.**
(**A**) Structures of A3Bctd (this study), A3A-ssDNA (pdb: 5sww), and A3Gctd-ssDNA (pdb: 6bux). Loop regions surrounding the active site and critical for substrate-binding are labelled. (**B**) Protein sequence alignment of A3Bctd, A3A, and A3Gctd. Identical residues are colored with blue boxes, and the loop 1, 3, and 7 regions are highlighted in yellow. Sequence alignment was done using Clustal Omega multiple sequence alignment tool.

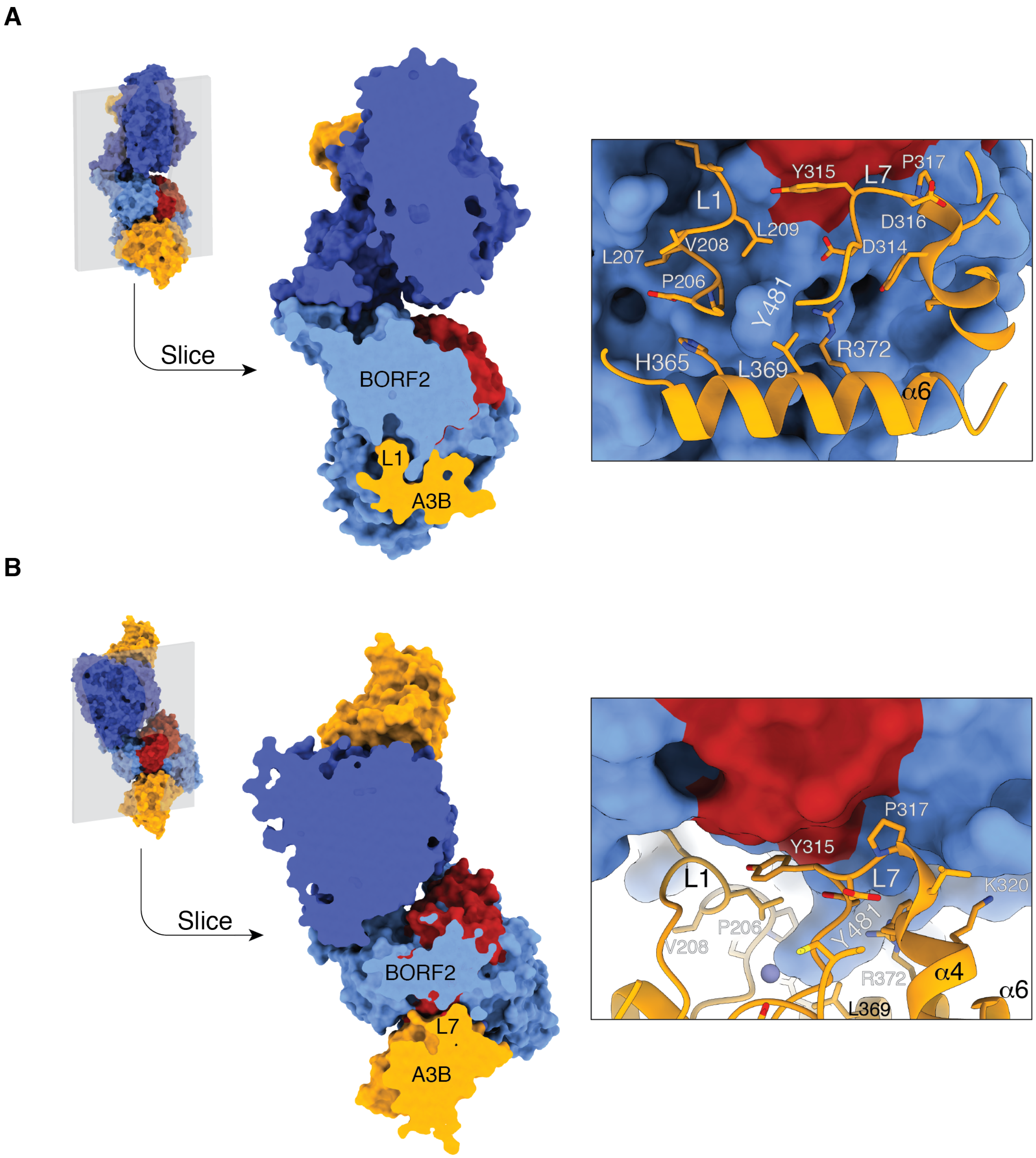


**Fig. S6. A3Bctd L1 and L7 interactions with BORF2.**

(**A**) A cross-sectional representation of the BORF2-A3Bctd complex providing additional details of the extensive interactions with L1. For instance, BORF2 residue 481 is inserted between L1 and helix-6 of A3Bctd.

(**B**) A different cross-sectional representation of the BORF2-A3Bctd complex providing additional details of the extensive interactions with L7. In particular, this view highlights the position of L7 relative to L1 and BORF2 SHI.

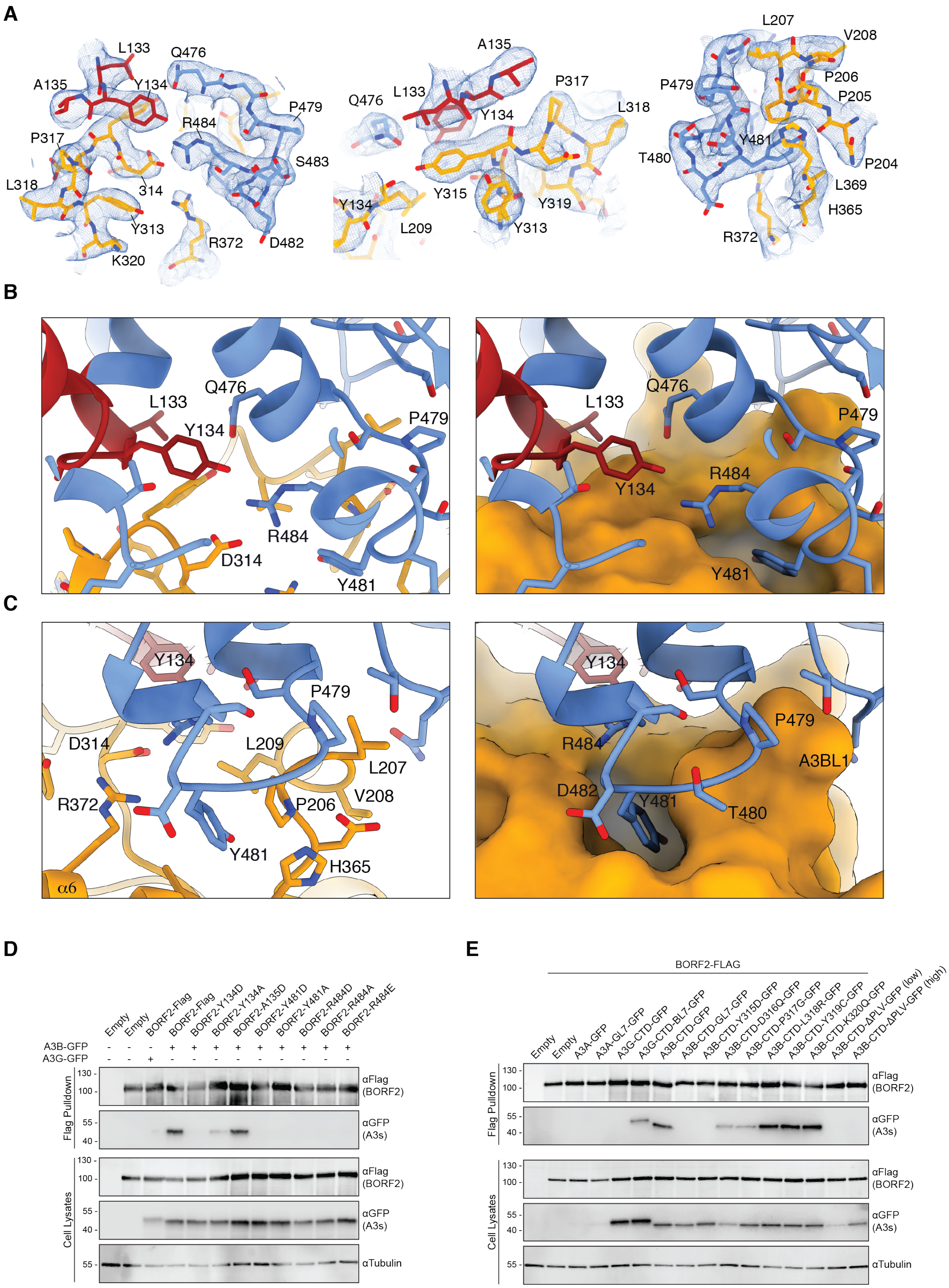
A
L133
Q476
A135
Y134
P479
R484
P317
S483
L318
314
Y313
R372
D482
K320
A135
Q476
L133
P317
L318
Y134
Y315
Y319
Y134
L209
Y313
L207
V208
P479
P206
P205
Y481
T480
P204
L369
H365
R372
B
L133
Q476
Y134
P479
R484
D314
Y481
C
Y134
P479
D314
L209
L207
R372
V208
P206
Y481
H365
α6
R484
A3BL1
D482
T480
D
Empty
BORF2-Flag
BORF2-Y134D
BORF2-Y134A
BORF2-A135D
BORF2-Y481D
BORF2-Y481A
BORF2-R484D
BORF2-R484A
BORF2-R484E
A3B-GFP
A3G-GFP
Flag Pulldown
Cell Lysates
αFlag (BORF2)
αGFP (A3s)
αTubulin
E
BORF2-FLAG
A3A-GFP
A3A-GL7-GFP
A3G-CTD-GFP
A3G-CTD-BL7-GFP
A3B-CTD-GFP
A3B-CTD-GL7-GFP
A3B-CTD-Y315D-GFP
A3B-CTD-D316Q-GFP
A3B-CTD-P317G-GFP
A3B-CTD-L318R-GFP
A3B-CTD-Y319C-GFP
A3B-CTD-K320Q-GFP
A3B-CTD-ΔPLV-GFP (low)
A3B-CTD-ΔPLV-GFP (high)

**Fig. S7. Summary of key interactions between BORF2 and A3Bctd.**

(**A**) Zoom-in of residues involved in interaction between BORF2 and A3Bctd with alternative views shown in the middle and right. BORF2 residues are colored blue and red, and A3Bctd residues are gold. Residues are depicted as sticks. Cryo-EM map represented by blue mesh.

(**B**) A zoom-in of the BORF2 SHI (blue/red) and A3Bctd L7 region (orange). Ribbon and stick schematic on the left, and the same view with a surface-filled representation of A3Bctd on the right.

(**C**) A zoom-in of BORF2 residue 481 (blue) and A3Bctd L1 region (orange). Ribbon and stick schematic on the left, and the same view with a surface-filled representation of A3Bctd on the right.

(**D**) Co-IP reactions with the indicated BORF2 constructs (anti-Flag) and A3Bctd-eGFP. A parallel reaction with BORF2 and A3Gctd-eGFP is included as a negative control.

(**E**) BORF2 (anti-Flag) co-IP experiments with the indicated A3-eGFP constructs including key A3Bctd mutants. The two A3A constructs were non-informative in this experiment but still included in the images here to prevent gel-cropping and allow visualization of the negative control reactions (empty +/- BORF2-flag).

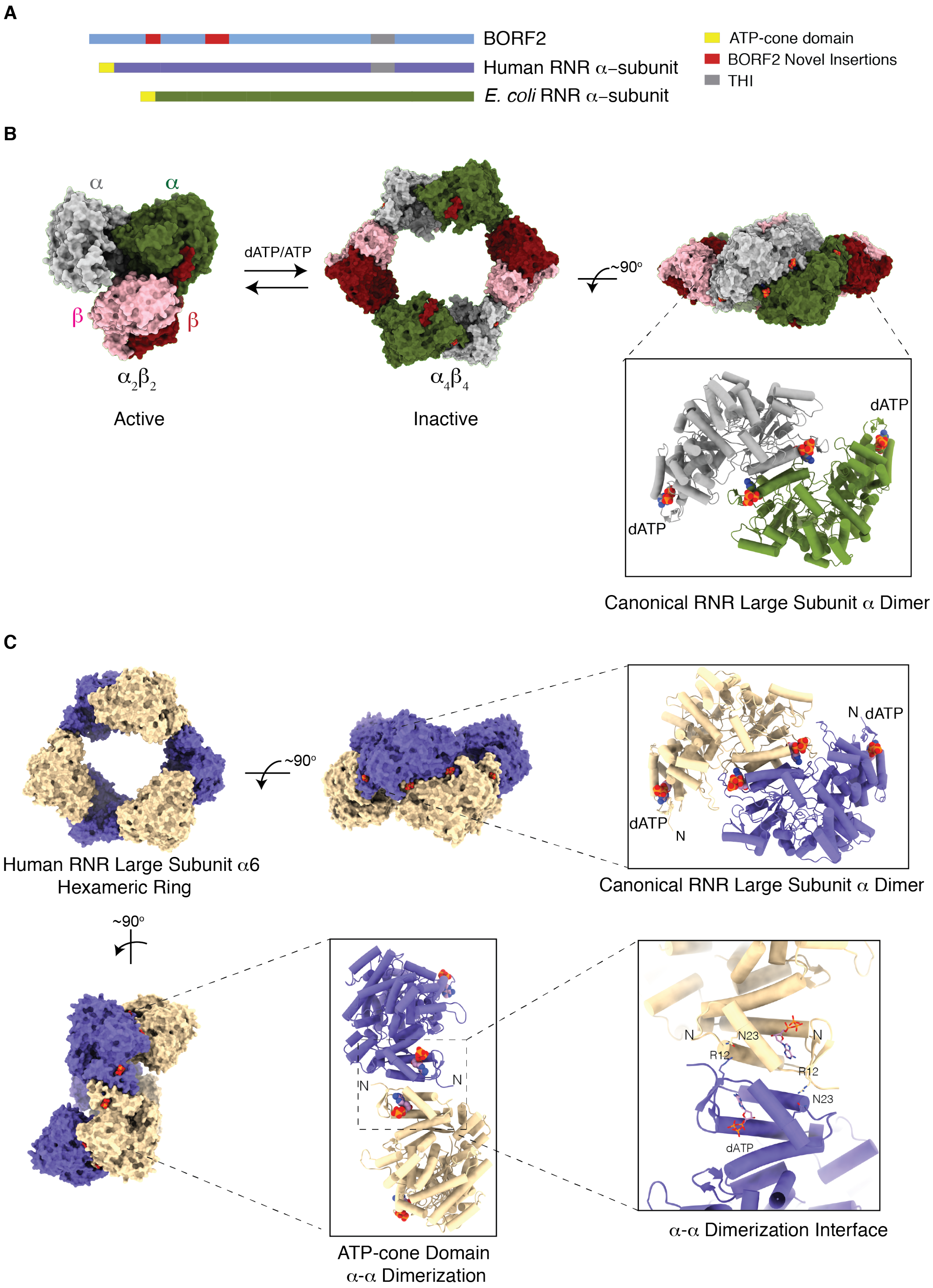
A
BORF2
Human RNR α–subunit
E. coli RNR α–subunit
ATP-cone domain
BORF2 Novel Insertions
THI
B
α
α
β
β
dATP/ATP
~90°
$\alpha_2\beta_2$
Active
$\alpha_4\beta_4$
Inactive
dATP
dATP
Canonical RNR Large Subunit α Dimer
C
~90°
N
dATP
dATP
N
Human RNR Large Subunit α6
Hexameric Ring
Canonical RNR Large Subunit α Dimer
~90°
N
N
N
N23
N
R12
R12
N23
dATP
ATP-cone Domain
α-α Dimerization
α-α Dimerization Interface

**Fig. S8. N-terminal ATP cone-domain is absent in EBV BORF2 despite key regulatory functions in other Class1a RNRs**

(**A**) Protein schematic of EBV BORF2, human RNR α-subunit, and *E. coli* RNR α-subunit with key domains indicated.

(**B**) Surface representations of the active and inactive forms of the *E. coli* RNR (pdb: 6w4x and pdb: 3UUS, respectively). The α-subunit is depicted in green and gray, and the β-subunit in pink and maroon. The active form is an $\alpha_2/\beta_2$ complex and dATP binding triggers the formation of an inactive $\alpha_4/\beta_4$ ring complex. The 90º rotation shows the canonical α/α dimer that exists in both complexes including dATP (spheres) bound to the cone domain.

(**C**) Surface representation of the dATP bound human RNR ring complex. This complex is comprised of two different dimeric forms of the RNR α domain - a canonical dimer (right) and a non-canonical dimer (below). The non-canonical dimer forms in part through the ATP-binding N-terminal region.

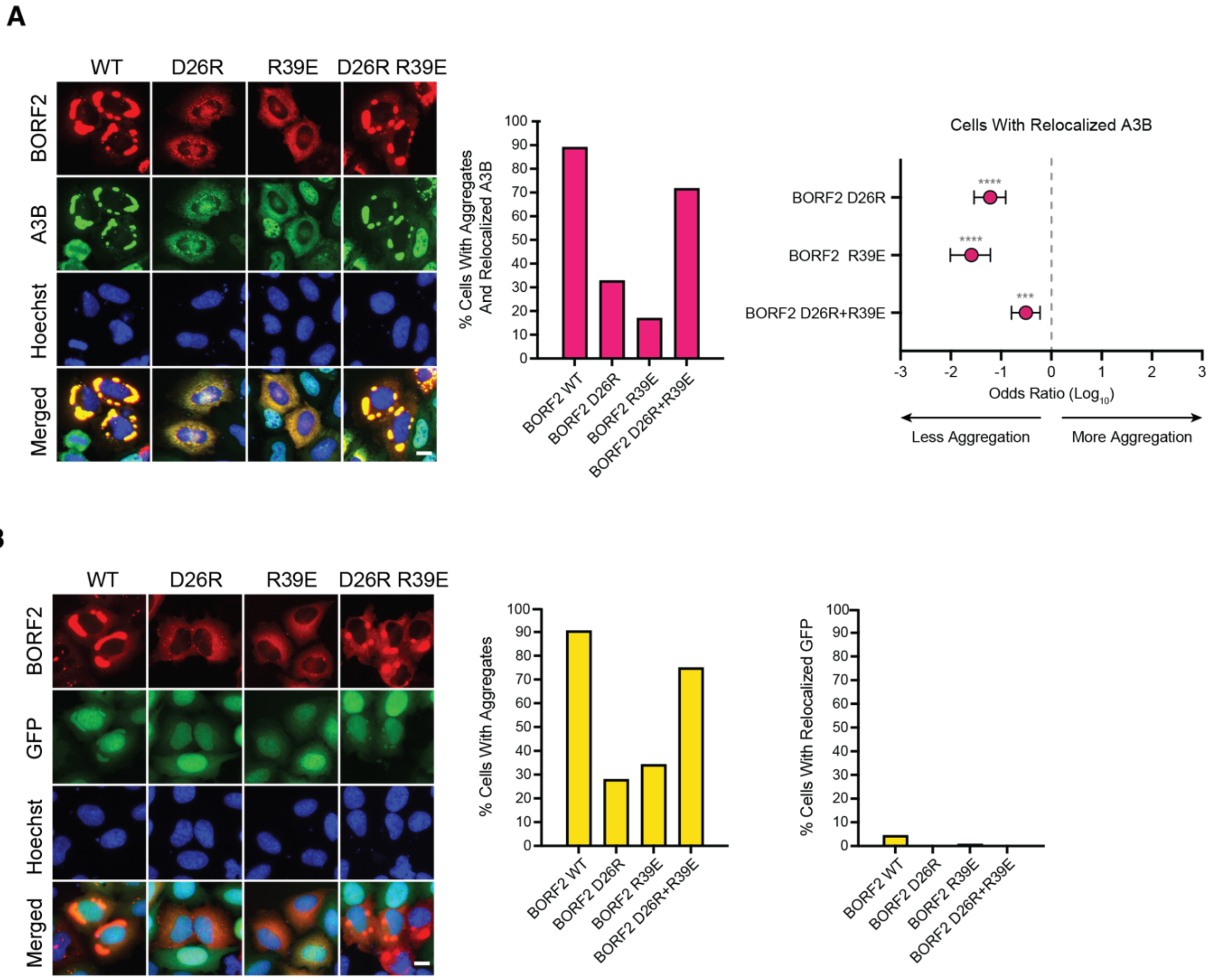


**Fig. S9. Cellular localization profile of BORF2 with full-length A3B.**

(**A**) Representative fluorescent microscopy images of A3B-eGFP and the indicated mCherry-BORF2 and constructs (scale = 10 µm). Nuclei are stained with Hoechst. Right: Quantification of aggregation and co-localization phenotypes (n > 100 for each condition). Odds ratios and p-values calculated by Fisher's exact test (ns, not significant; *** $P \leq 0.001$; **** $P \leq 0.0001$). Error bars = 95% confidence intervals.

(**B**) Representative fluorescent microscopy images of eGFP and the indicated mCherry-BORF2 constructs (scale = 10 µm). Nuclei are stained with Hoechst. Right: Quantification of aggregation and co-localization phenotypes ($n \geq 100$ for each condition). This control was included to show that the BORF2 aggregates occur in the absence of A3B but are large enough to trap some eGFP. The quantification histogram at the lower right indicates that eGFP trapping (presumably passive) occurs at much lower levels than A3Bctd-eGFP (Fig. 3C-D) or full-length A3B-eGFP (panel A) co-localization with the BORF2 aggregates (active and highly selective as shown here).

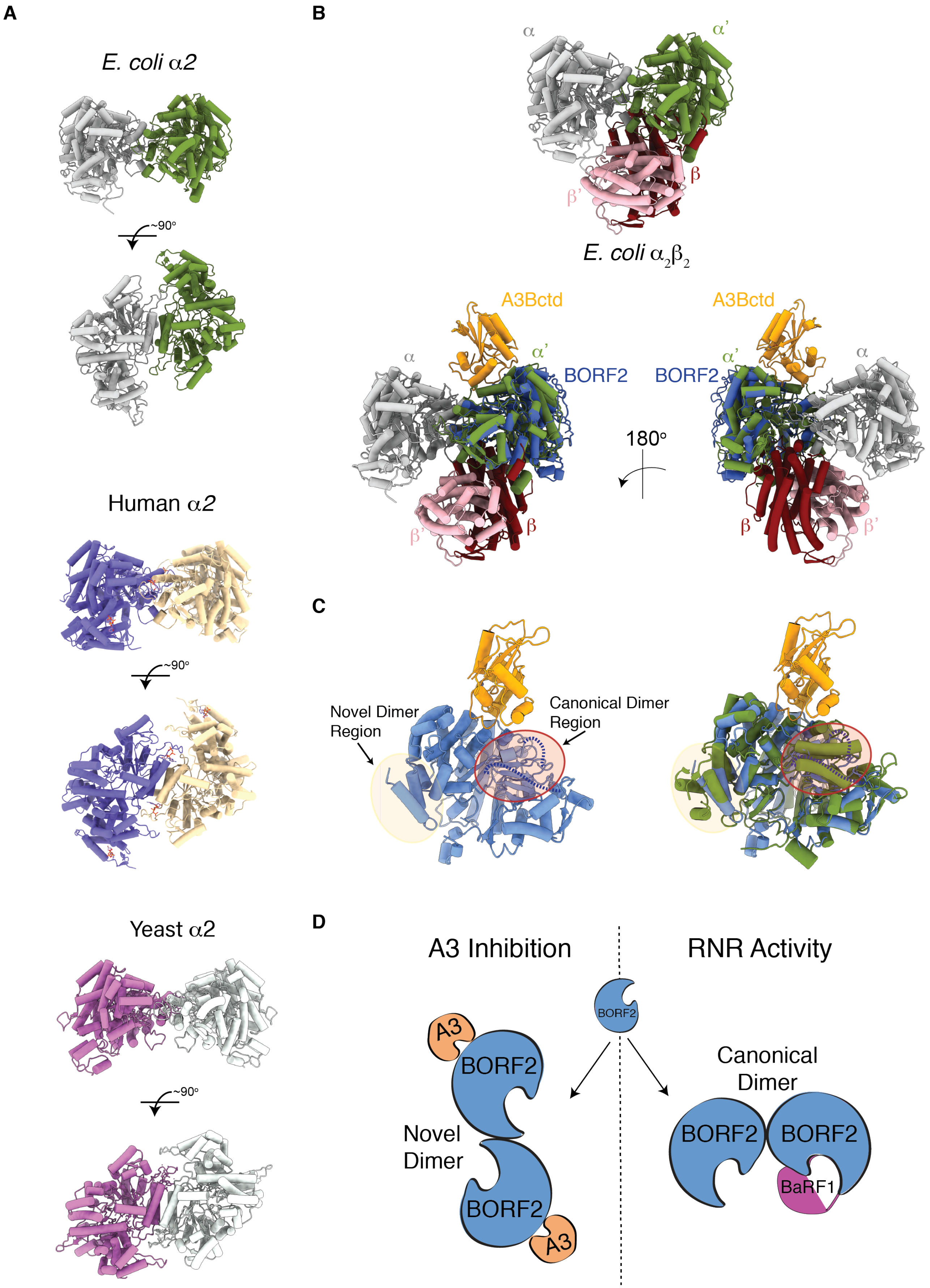
A
E. coli α2
~90°
~90°
Human α2
~90°
Yeast α2
B
α
α'
β
β'
E. coli $\alpha_2\beta_2$
A3Bctd
A3Bctd
α
α'
BORF2
BORF2
α'
α
180°
β'
β
β
β'
C
Novel Dimer Region
Canonical Dimer Region
D
A3 Inhibition
RNR Activity
BORF2
A3
BORF2
Novel Dimer
BORF2
A3
Canonical Dimer
BORF2
BORF2
BaRF1

**Fig. S10. Models for BORF2 function in ribonucleotide reduction and A3B inhibition.**

(**A**) Ribbon schematics of *E. coli* (pdb: 6w4x), human (pdb: 6aui), and yeast RNR (pdb: 2eud) canonical $\alpha$-subunit dimers.

(**B**) Top - ribbon schematic of *E. coli* $\alpha2/\beta2$ structure (pdb: 6w4x). Bottom – overlay the *E. coli* structure with a single BORF2-A3Bctd complex. The predicted BaRF1 binding surface based on the positioning of the *E. coli* $\beta$-subunit is distinct from the observed A3B binding surface.

(**C**) Left - ribbon schematic of BORF2-A3B monomer with a yellow circle showing the BORF2 dimer region and a red circle highlighting the predicted canonical dimerization domain. Blue dashed lines represent BORF2 disordered regions. Right – overlay of BORF2-A3Bctd with the *E. coli* $\alpha$ subunit (pdb: 6w4x). Yellow circle showing BORF2 dimer region. Red circle showing the *E. coli* $\alpha$ subunit dimerization domain that is disordered in the BORF2-A3Bctd structure. Blue dashed lines represent BORF2 disordered regions.

(**D**) Cartoon schematic of the two different BORF2 dimerization mechanisms, one observed here for A3B inhibition (left) and the other predicted to be required for ribonucleotide reduction (right).

**Movie S1. 3D Variability analysis of BORF2-A3Bctd complex.** Top view of the complex. This analysis was performed in cryoSPARC using the 3D variability feature.

**Movie S2. 3D Variability analysis of BORF2-A3Bctd complex.** Side view of the complex. This analysis was performed in cryoSPARC using the 3D variability feature.

**Data collection and processing**

| | | | |
|---|---|---|---|
| Microscope | | FEI Titan Krios | |
| Voltage (kV) | | 300 | |
| Detector | | Gatan K3 | |
| Energy filter width (eV) | | 20 | |
| Magnification (nominal X) | | 81,000 | |
| Pixel size (Å) | | 0.844 (0.422 super-resolution) | |
| Defocus range (µm) | | 0.8-2.5 | |
| Total Dose (e-/ Å$^2$) | | 60 | |
| Number of frames | | 50 | |
| Exposure time/movie (sec) | | 5.22 | |
| Total Micrographs collected | | 7,488 | |
| Micrographs used | | 6,759 | |
| Automation Software | | SerialEM | |
| Particles extracted (total) | | 7,675,126 | |
| | **Overall Global Resolution** | **BORF2-A3Bctd** Focused-refinement around BORF2-A3B monomer | **Composite** Generated from focused refined map using Phenix: combine focused maps |
| EMD # | EMD-24716 | EMD-24715 | EMD-24709 |
| PDB # | | | 7wr6 |
| Particles | 121,596 | 243,192 (Symmetry Expansion) | |
| Symmetry Imposed | C2 | C1 | |
| Map Sharpening B factor | -91.5 | n/a DeepEMhancer | n/a DeepEMhancer |
| 0.5 FSC (Unmasked/Masked) (Å) | 6.03/3.33 | 4.31/3.01 | |
| 0.143 FSC (Unmasked/Masked) (Å) | 3.84/2.82 | 3.6/2.55 | |

**Refinement and validation**

| | | | |
|---|---|---|---|
| Refinement package | | | Phenix |
| Refinement tool* | | | Real-space refine |
| Model composition | | | |
| | | | 13,130 |

| | |
|---|---|
| Non-hydrogen atoms | 1634 |
| Protein residues | 2 |
| Ligands ($Zn^{2+}$) | |
| B factors (Å$^2$) | |
| Protein | 71.81 |
| Ligand | 95.38 |
| R.m.s deviations | |
| Bond lengths (Å$^2$) | 0.003 |
| Bond angles (º) | 0.570 |
| Ramachandran Plot | |
| Favored (%) | 94.21 |
| Allowed (%) | 5.79 |
| Disallowed (%) | 0 |
| Validation | |
| MolProbity score | 1.88 |
| Clashscore | 9.11 |
| Poor rotomers (%) | 0 |
| C-beta outliers (%) | 0 |
| CaBLAM outliers (%) | 0.25 |
| CC (mask) | 0.8 |
| EMRinger | 4.10 |

**Table S1. Cryo-EM data collection and structure refinement data**
